# Gene replacement therapy for Lafora disease in the *Epm2a^-/-^* mouse model

**DOI:** 10.1101/2023.12.14.571636

**Authors:** Luis Zafra-Puerta, Daniel F. Burgos, Nerea Iglesias-Cabeza, Juan González-Fernández, Gema Sánchez-Martín, Marina P. Sánchez, José M. Serratosa

**Affiliations:** Laboratory of Neurology, Instituto de Investigación Sanitaria-Fundación Jiménez Díaz, Universidad Autónoma de Madrid (IIS-FJD, UAM), 28040 Madrid, Spain; Centro de Investigación Biomédica en Red de Enfermedades Raras (CIBERER), 28029 Madrid, Spain; PhD Program in Neuroscience, Universidad Autonoma de Madrid-Cajal Institute, Madrid, Spain 28029; Fondazione Malattie Rare Mauro Baschirotto BIRD Onlus, Longare (VI), Italy; Departament of Microbiology and Parasitology, Faculty of Pharmacy, Complutense University of Madrid

**Keywords:** Progressive myoclonic epilepsy, *Epm2a^-/-^* knock-out mouse, Gene therapy, rAAV, Polyglucosans, Laforin

## Abstract

Lafora disease is a rare and fatal form of progressive myoclonic epilepsy typically occurring early in adolescence. Common symptoms include seizures, dementia, and a progressive neurological decline leading to death within 5-15 years from onset. The disease results from mutations transmitted with autosomal recessive inheritance in the *EPM2A* gene, encoding laforin, a dual-specificity phosphatase, or the *EPM2B* gene, encoding malin, an E3-ubiquitin ligase. Laforin has glucan phosphatase activity, is an adapter of enzymes involved in glycogen metabolism, is involved in endoplasmic reticulum-stress and protein clearance, and acts as a tumor suppressor protein. Laforin and malin work together in a complex to control glycogen synthesis and prevent the toxicity produced by misfolded proteins via the ubiquitin-proteasome system. Disruptions in either protein can lead to alterations in this complex, leading to the formation of Lafora bodies that contain abnormal, insoluble, and hyperphosphorylated forms of glycogen called polyglucosans.

We used the *Epm2a^-/-^* knock-out mouse model of Lafora disease to apply a gene replacement therapy by administering intracerebroventricular injections of a recombinant adeno-associated virus carrying the human *EPM2A* gene. We evaluated the effects of this treatment by means of neuropathological studies, behavioral tests, video-electroencephalography recording, and proteomic/phosphoproteomic analysis.

Gene therapy with recombinant adeno-associated virus containing the *EPM2A* gene ameliorated neurological and histopathological alterations, reduced epileptic activity and neuronal hyperexcitability, and decreased the formation of Lafora bodies. Differential quantitative proteomics and phosphoproteomics revealed beneficial changes in various molecular pathways altered in Lafora disease. Improvements were observed for up to nine months following a single intracerebroventricular injection. In conclusion, gene replacement therapy with human *EPM2A* gene in the *Epm2a^-/-^* knock-out mice shows promise as a potential treatment for Lafora disease.

## Introduction

Lafora disease (OMIM #254780; ORPHA: 501) is a rare and fatal form of neurogenerative disease that appears in early adolescence with seizures and a progressive neurological decline with dementia resulting in death within 5-15 years^1–3^. Unfortunately, there is no specific therapy, and patients can only receive antiseizure medications to temporarily manage seizures^4^. The disease is caused by autosomal recessive inherited mutations in either the *EPM2A* (OMIM 607566) gene, encoding the dual-specificity phosphatase laforin^5–8^, or the *EPM2B* (OMIM 608072) gene, encoding the E3-ubiquitin ligase malin^9,10^. Laforin is a glucan phosphatase acting as an adapter protein of enzymes involved in glycogen synthesis, an adapter protein in endoplasmic reticulum (ER)-stress and protein clearance, and a tumor suppressor protein^11^. The laforin-malin complex regulates glycogen synthesis by inducing proteasome-dependent degradation of muscle glycogen synthase (GS), glycogen debranching enzyme (GDE), and protein targeting to glycogen (PTG)^11,12^. Additionally, the laforin-malin complex helps mitigate the toxicity produced by misfolded proteins through the ubiquitin-proteasome system (UPS) ^13^. Disruptions in laforin or malin lead to the formation of aggregates of abnormal, insoluble, poorly branched and hyperphosphorylated forms of glycogen, known as Lafora bodies (LB)^14–17^. Alterations in oxidative stress, protein misfolding, and proteasomal dysfunction, also contribute to the pathophysiology of the disease^18–20^.

Different murine models of Lafora disease have been generated, including the *Epm2a^-/-^* and *Epm2b^-/-^* knock-out mice^21,22^. These models replicate, although with a milder phenotype, most of the neurological alterations seen in patients^23^. *Epm2a^-/-^* and *Epm2b^-/-^* knock-out mouse models have been used to assay different putative treatments to cure or ameliorate the symptoms of the disease. Thus, we showed in these models that metformin improved many neurological alterations^24,25^ Subsequently, we performed a study with metformin treatment in early-stage patients and showed that it slowed symptom progression, resulting in a slower decline in daily life activities^24^ compared to treatments in patients in more advanced stages of the disease^26^. Alternative therapeutic strategies, such us sodium selenate^27^, VAL-0417^28^, the antisense oligonucleotide (ASO) *Gys1* (to target GS)^29^ and various modulators of neuroinflammation have also been assessed in animal models^30^. Adeno-associated (AAV)-based downregulation of the *Gys1* gene through CRISPR-CAS9^31^ or miRNA^32^ also resulted in a remarkable reduction in polyglucosan body formation and a slight reduction in some neuroinflammatory markers.

Recombinant AAV (rAAV) are widely used vectors for gene replacement therapy. They are non-pathogenic viruses that produce low immunological responses, rarely integrate into the host genome, have a broad tropism, and allow long-term transgene expression. These viruses are small, non-enveloped, single-stranded (ss) DNA particles belonging to the *Parvoviridae* family, genus *Dependoviridae*. To date, more than 10 different AAV serotypes and more than 100 variants have been isolated from adenovirus stocks and human/non-human primate tissues. Different serotypes have different tropism and recent progress in the field has led to precise targeting towards specific tissues of interest^33–41^.

Here, we show that a new rAAV2/9-CAG-h*EPM2A* (rAAV-h*EPM2A*) vector, containing the human *EPM2A* (h*EPM2A*) gene, significantly diminishes neurological and histopathological alterations, reduces epileptic activity, and decreases LB formation in the *Epm2a^-/-^* mouse model of Lafora disease. Additionally, we show through differential quantitative proteomic and phosphoproteomic analysis that human laforin produces beneficial changes in certain molecular pathways altered in Lafora disease. This molecular analysis is crucial for comprehending the gene therapy mechanisms and understanding the specific processes that are being corrected by *EPM2A* gene replacement.

## Materials and methods

### Experimental Animals

We used the *Epm2a*^-/-^ mouse model of Lafora disease, generated following previously described methods^21^, and age-matched wild-type (WT) mice (C57BL6). Since no-gender related phenotype differences have been described in mice^42^ or patients with Lafora disease^26,43^, we analyzed data from male and female mice indistinctively. The mouse colonies were bred in the Animal Facility Service of the Instituto de Investigación Sanitaria-Fundación Jiménez Díaz and were housed in isolated cages with a 12:12 light/dark cycle at a constant temperature of 23 °C, with free access to food and water. All experiments were conducted with the utmost care to use and sacrifice the minimum number of animals while minimizing their suffering. The experimental procedures adhered to the "Principles of Laboratory Animal Care" (NIH publication No. 86–23, revised 1985), as well as the European Communities Council Directive (2010/63/EU) and were approved by the Ethical Review Board of the Instituto de Investigación Sanitaria-Fundación Jiménez Díaz.

### Production of rAAV2/9-CAG-h*EPM2A*, rAAV2/9-CAG-*GFP* and rAAV2/9-CAG-Null vectors

The rAAV2/9-CAG-h*EPM2A* (rAAV-h*EPM2A*) vector, containing the cDNA of the *EPM2A* gene transcript variant 1 (hE*PM2A)* (NM_005670.4), the rAAV2/9-CAG-Null (rAAV-Null) vector, containing a non-coding DNA, and the rAAV2/9-CAG-*GFP* (rAAV-*GFP*) vector, containing the green fluorescent protein (*GFP*) gene, were generated in the Unitat de Producció de Vectors (UPV_ www.viralvector.eu). The production of those vectors was performed following the triple transfection system: (a) the ITR-containing plasmid; (b) the plasmid encoding AAV capsid (VP1, VP2 and VP3 proteins) and replicate genes; and (c) the adenoviral helper plasmid. To remove empty capsids, AAV vectors were purified by iodixanol-based ultracentrifugation^44^.

### Stereotaxic intracerebroventricular (ICV) injections

Three-month-old *Epm2a^-/-^* mice received a single ICV injection of rAAV-h*EPM2A*, rAAV-*GFP* or rAAV-Null vectors. Mice were anesthetized in an induction chamber filled with 4% isoflurane and 2% O^2^ and maintained with 2% isoflurane and 1.5% O_2_. Mice were fixed in a stereotaxic frame (Stoelting, Illinois, USA) and body temperature was maintained using a heating pad at 37°C. Hydration was controlled with a subcutaneous saline injection (1 mL), and ophthalmic gel was applied to prevent dry eyes. The total number of mice subjected to rAAV-h*EPM2A* and rAAV-Null ICV injections was 25-35 per group and condition.

The incision site was sterilized with 70% ethanol, and it was made in the midline, starting behind the eyes. Bregma and lambda were shown with H_2_O_2_ on the skull. A small burr hole was drilled according to stereotaxic coordinates of the right cerebral lateral ventricle relative to bregma (anterioposterior -0.3 and mediolateral -0.9). A Hamilton^®^ syringe (ThermoFisher Scientific, Massachusetts, USA, Cat. #10664301) was introduced at -2.5 cm in the dorsoventral axis to deliver 3 µL of viral suspension with a titer of 1.26 x 10^12^ vg/mL at a rate of 3 µL/min. The incision was sutured, and meloxicam (Boehringer Ingelheim, Georgia, USA) (5mg/kg) was administered as an analgesic.

### RNA extraction and quantitative reverse transcription-polymerase chain reaction (RT-qPCR)

RNA was extracted from brain samples previously homogenized on ice with TRIzol^TM^ Reagent (ThermoFisher Scientific, Massachusetts, US). The RNA pellets were washed, dried, resuspended, and treated with DNase Enzyme (ThermoFisher Scientific, Massachusetts, US), and quantified with a NanoDrop ND-1000 spectrophotometer (ThermoFisher, Massachusetts, US). RT-PCR experiments were carried out using the High-Capacity cDNA Reverse Transcription Kit with RNase inhibitor (ThermoFisher Scientific, Massachusetts, US) with 1 μg of RNA per reaction. The RT-PCR conditions were 25°C for 10 min – 37°C for 120 min – 85°C for 5 min. For RT-qPCR, cDNA from the h*EPM2A* transcript variant 1 was used as a template. The reaction was performed with TaqMan™ Fast Advanced Master Mix (ThermoFisher Scientific, MA, USA), with *Epm2a*, *EPM2A* and *Gapdh* probes (ThermoFisher Scientific, MA, USA). The qPCR conditions were 50°C for 2 min – 95°C for 2 min – 95°C for 1 sec and 60°C for 20 sec (40 cycles). Analysis was performed using the 2^-ΔΔCT^ method.

### Immunofluorescence, immunohistochemistry and PAS-diastase staining

Immunohistochemistry (IHC) and PAS-diastase (PAS-D) staining procedures were performed as previously described^45^ and in Supplementary Material. For IHC, the primary antibodies used were green fluorescent protein (GFP) (1:100 dilution; Abcam, Cambridge, UK; Cat. #ab183734), neuronal nuclei (NeuN) (1:100 dilution; Millipore, Temecula, CA, USA; Cat. # MAB377) and laforin (3µg/mL; Lifespan Biosciences, Washington, US, Cat. #LS B6474). For IF-P, the primary antibody was glial fibrillary acidic protein (GFAP) (1:1000 dilution; Millipore, Temecula, CA, USA; Cat. #MAB360). Secondary antibodies were conjugated to Alexa Fluor 594 (donkey anti-mouse, 1:400 dilution; Abcam, Cambridge, UK, Cat. #ab150108). Samples from 4-6 mice per group were used, and two consecutive sections per animal were stained and analyzed. Images from the same area of the CA1 region of each hippocampus were acquired using a Leica DMLB 2 microscope (Leica, Wetzlar, Germany) connected to a Leica DFC320 FireWire digital microscope camera (Leica, Wetzlar, Germany) for IHC processed sections, and with a Zeiss Axioscope 5 (Zeiss, Jena, Germany) connected to an Axiocam 208 color camera (Zeiss, Jena, Germany) for sections processed via IF-P. Subsequently, LBs and NeuN- or GFAP-positive cells were quantified by two researchers using ImageJ software (NIH, Bethesda, MD, USA). Reported values represents the average of these quantifications.

### Spontaneous locomotor activity

Spontaneous movements were monitored with a computerized actimeter (Harvard Apparatus, Holliston, MA, USA) and with the SEDACOM 1.4 software (Harvard Apparatus, Holliston, MA, USA) as described previously^45^ and in Supplementary Material.

### Motor coordination

Motor coordination and balance were assessed using the rotarod test (Harvard Apparatus, Holliston, MA, USA). Detailed procedures of these analyses are explained in^45^ and in Supplementary Material.

### Object recognition task

The object recognition task (ORT) was used to assess episodic memory retention, as reported in^45^ and in Supplementary Material.

### Sensitivity to Pentylenetetrazole (PTZ)

To analyze neuronal hyperexcitability, PTZ was administered via intraperitoneal (IP) injection, as explained in^45^ and in Supplementary Material.

### Video-EEG analysis

A plastic pedestal (Plastics1, Virginia, USA) with trimmed electrodes was surgically implanted and secured with acrylic resin onto the skull. Post-surgical pain was managed with meloxicam (5 mg/kg) (Boehringer Ingelheim, GA, USA). Animals were allowed one week for recovery before testing. Video-EEG recordings were obtained using a wireless transmitter (Epoch, CA, USA) attached to the pedestal, and the data were digitally recorded on a computer under free-motion conditions. More details on the video-EEG analyses are described in^45^ and in the Supplementary Material.

### Differential quantitative proteomics by isobaric labeling: TMT11plex

For differential quantitation in the proteomic and phosphoproteomic analysis, peptides were labeled with the Tandem Mass Tag 11 Plex (TMT11-Plex) technique. Identification and quantification of proteins were performed using liquid chromatography coupled with tandem mass spectrometry (LC-MS/MS)^46,47^.

Data were analyzed using a logarithmic statistical model (log2), enabling the estimation of peptide and protein abundances (Zq). For the quantification of changes in peptide phosphorylation, the abundance of phospho-peptides was assessed using the corresponding standardized log2 ratio (Zp)^46,47^.

### Statistical analysis

Values are presented as mean ± standard error of the mean (SEM) or as percentages. To analyze the differences between experimental groups, we employed one- or two-way ANOVA, Fisher’s exact test and non-parametric Kruskal-Wallis or Mann-Whitney test, as indicated in each specific case. For EEG analysis, the Area Under the Curve (AUC) was obtained to compare the differences in the power spectra between groups. For proteomics and phosphoproteomics, changes in protein and phospho-peptides abundance were determined by comparing the means of Zq and Zp, respectively, between groups and selecting those with a *p-*value of 0.05 or lower. Statistical analyses were conducted using GraphPad Prism 8.0 (San Diego, CA). Statistical tests were two-tailed and the statistical significance thresholds were denoted as *p < 0.05, **p < 0.01, ***p < 0.001, and ****p < 0.0001.

## Data availability statement

Data supporting the findings of this study are available from the corresponding author.

## Results

### The rAAV-h*EPM2A* vector efficiently transduces central nervous system (CNS) cells in *Epm2a^-/-^* mice, leading to transcription and translation of the h*EPM2A* transgene within the CNS

Three months after a single ICV injection of rAAV-*GFP* or rAAV-h*EPM2A* in 3-month-old *Epm2a^-/-^* mice, we verified effective transduction in CNS cells. Firstly, we confirmed proper h*EPM2A* transgene transcription through RT-PCR **(Supplementary Figure S1A).** Next, with IHC using an antibody against GFP, we observed widespread distribution of the vector throughout the CNS **(Supplementary Figure S1B).**

Subsequently, transcription of the h*EPM2A* transgene was quantified by RT-qPCR (**Fig. 1A-B)**. The h*EPM2A* transcript levels were 60 and 80% of those observed in WT animals at 6 and 12 months of age, respectively **(Fig. 1A-B)**. Additionally, RNA translation was analyzed by IHC with an antibody against laforin, demonstrating successful protein expression in the hippocampus, cortex, and areas around the lateral ventricles, 3 and 9 months after the ICV injection of rAAV-h*EPM2A* **(Fig. 1C)**.

**Figure 1.**
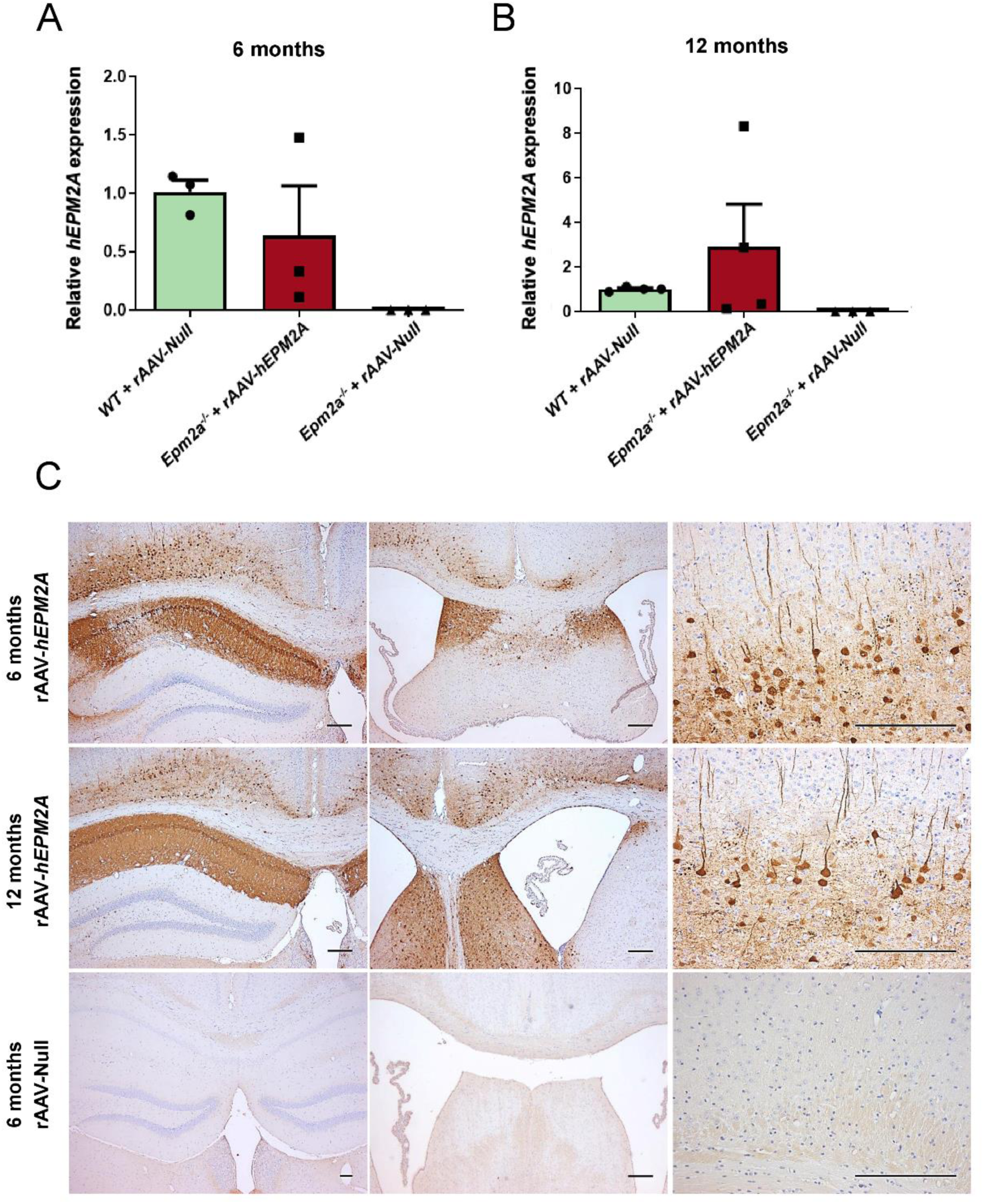
Quantification of h*EPM2A* transgene expression and distribution of human laforin after a single ICV injection of rAAV-h*EPM2A*. (**A-B**) RT-qPCR quantification showing *Epm2a* and *EPM2A* cDNA levels in brain samples obtained from WT and *Epm2a^-/-^* mice 3 months (**A**) or 9 months (**B**) post-ICV injection of either rAAV-h*EPM2A* or rAAV-Null. (**C**) IHC with an anti-laforin antibody showing the distribution of human laforin protein across the hippocampus, areas surrounding the lateral ventricles, and cortical neurons. n= 4-6 mice per group and experiment. Scale bar =100µm

### ICV injection of rAAV-h*EPM2A* prevents LB formation in the brain of treated *Epm2a^-/-^* mice

We analyzed LBs in the CA1 region of the hippocampus with PAS-D staining 3 and 9 months following ICV injections of rAAV-h*EPM2A* and rAAV-Null vector (**Fig. 2A)**. We also quantified the progression of LB formation from 3-month-old *Epm2a^-/-^* mice without treatment, and at 3 and 9 months post-ICV injection **(Fig. 2A-B)**. Mice treated with rAAV-h*EPM2A* showed fewer LBs **(Fig. 2A)** and a slower progression of LB formation compared to animals injected with rAAV-Null **(Fig. 2B).**

**Figure 2.**
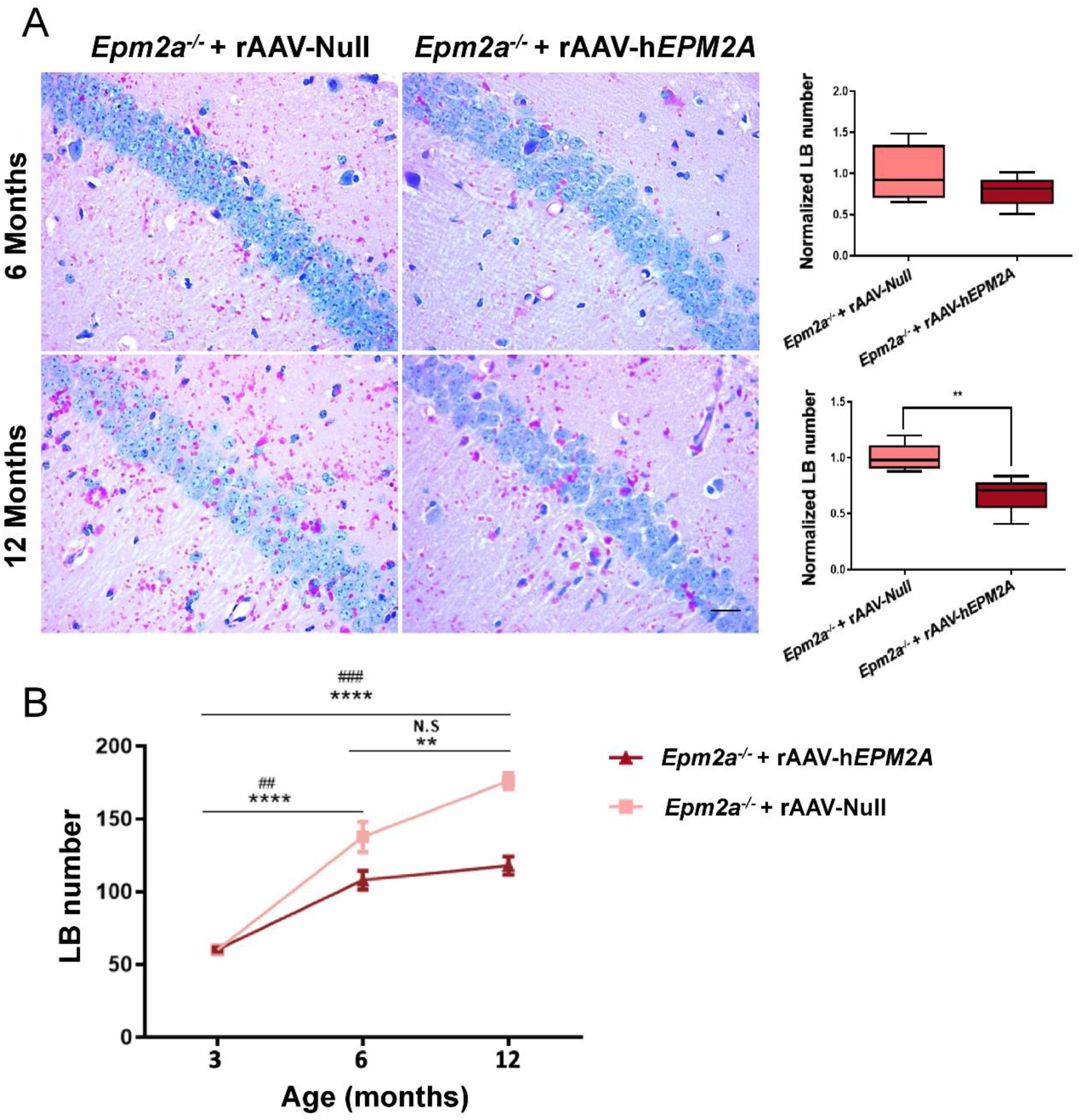
Quantification of LB formation in the CA1 region of the hippocampus of rAAV-h*EPM2A* treated *Epm2a^-/-^* mice. (**A**) PAS-D staining in *Epm2a^-/-^* mice 3 and 9 months after ICV injection of rAAV-h*EPM2A* or rAAV-Null and quantitative comparison of LB number. Results are expressed as median of independent samples. Whiskers in box plots show the minimum and maximum values. Means were normalized using *Epm2a^-/-^* mice injected with rAAV-Null values. (**B**) Comparative progression of LB formation over time in *Epm2a^-/-^* mice treated with rAAV-h*EPM2A* or rAAV-Null. A non-parametric Mann-Whitney test and two-way ANOVA test with Tukey’s multiple comparisons were performed. * p < 0.05, **** p < 0.0001. Symbols indicate: * for differences in LB progression over time in *Epm2a^-/-^* mice injected with rAAV-Null; and # for differences in LB progression over time in *Epm2a^-/-^* mice treated with rAAV-h*EPM2A*. n = 4-6 mice per group. Scale bar =25µm

### Treatment with rAAV-h*EPM2A* in *Epm2a^-/-^*mice decreases astrogliosis but not neurodegeneration

The effect of rAAV-h*EPM2A* on neuroinflammation in *Epm2a^-/-^* mice was assessed 3 and 9 months after ICV injections, using a GFAP antibody **(Fig. 3A)**. Quantification of GFAP-positive cells present in the CA1 region of the hippocampus of 6-**(Fig. 3B)** and 12-month-old **(Fig. 3C)** *Epm2a^−/−^* mice showed fewer GFAP-immunostained cells in the brains of rAAV-h*EPM2A* treated mice compared to those injected with the null vector.

**Figure 3.**
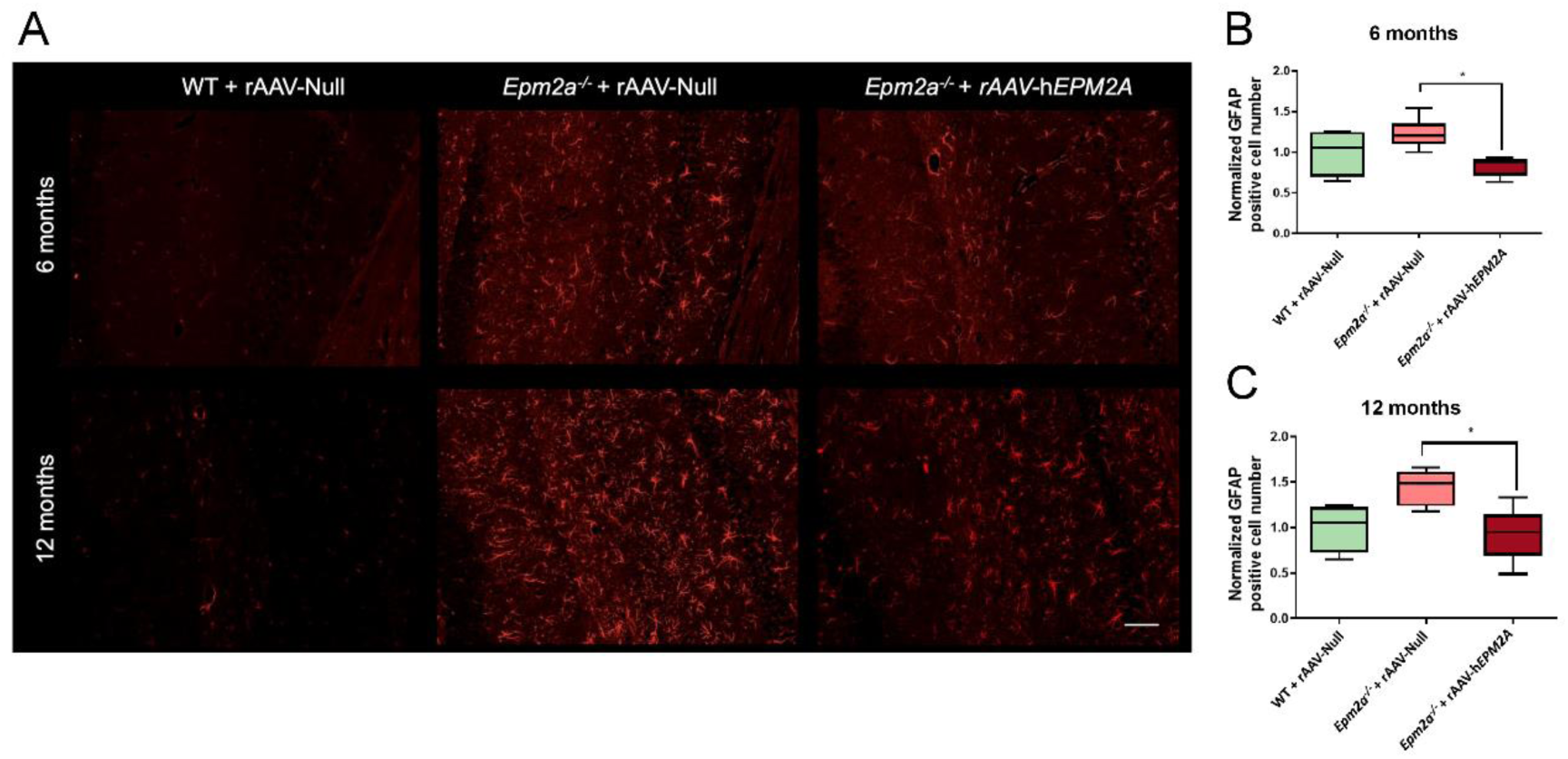
Neuroinflammation in the CA1 region of the hippocampus of treated *Epm2a^-/-^* mice. (**A**) IF-P with an anti-GFAP antibody staining reactive astrocytes. (**B-C**) Quantification of reactive astrocytes in the hippocampal CA1 area of WT and *Epm2a^-/-^* mice treated with rAAV-h*EPM2A* or rAAV-Null 3 months (**B**) or 9 months (**C**) after ICV injection. Results are expressed as median of independent samples. Whiskers in box plots show the minimum and maximum values. Means were normalized using WT mice injected with rAAV-Null values. A non-parametric Kruskal-Wallis test was performed followed by Dunńs multiple comparisons. * p < 0.05. n = 4-6 mice per group. Scale bar =100µm

Additionally, the effect of rAAV-h*EPM2A* on neurodegeneration was examined using NeuN antibody at 3 and 9 months post-ICV injection (**Supplementary Fig. S2A)**. A slight, albeit statistically non-significant, reduction in neurodegeneration was observed in the CA1 of 6-month-old mice treated with rAAV-h*EPM2A* in comparison to aged-matched *Epm2a^-/-^* mice injected with rAAV-Null (**Supplementary Fig. S2B)**. However, this trend was not evident 9 months after ICV injections (**Supplementary Fig. S2C).**

### Gene therapy with rAAV-h*EPM2A* ICV injection in *Epm2a^-/-^* mice delays the onset of memory decline and diminishes motor impairments

We analyzed the effects of laforin expression on episodic memory, motor coordination, and spontaneous locomotor activity. Memory performance was significantly improved in 6-month-old *Epm2a^-/-^* treated mice, as indicated by their higher discrimination index **(Fig. 4A)**. This improvement, however, was not evident in 12-month-old *Epm2a^-/-^* treated mice **(Fig. 4B)**, suggesting that rAAV-h*EPM2A* delayed the onset of spatial memory deficits but not in a sustained manner over time. Motor coordination was also improved with treatment, as evidenced by an increased latency to fall from the rod in the rotarod test at both 3 and 9 months post-ICV injection (**Fig. 4C-D**). Analysis of spontaneous motor activity in the actimeter revealed enhanced performance of *Epm2a^-/-^* treated mice 3 months after ICV injection **(Fig. 4E).** However, these improvements were not maintained 9 months after ICV injection **(Fig. 4F)**.

**Figure 4.**
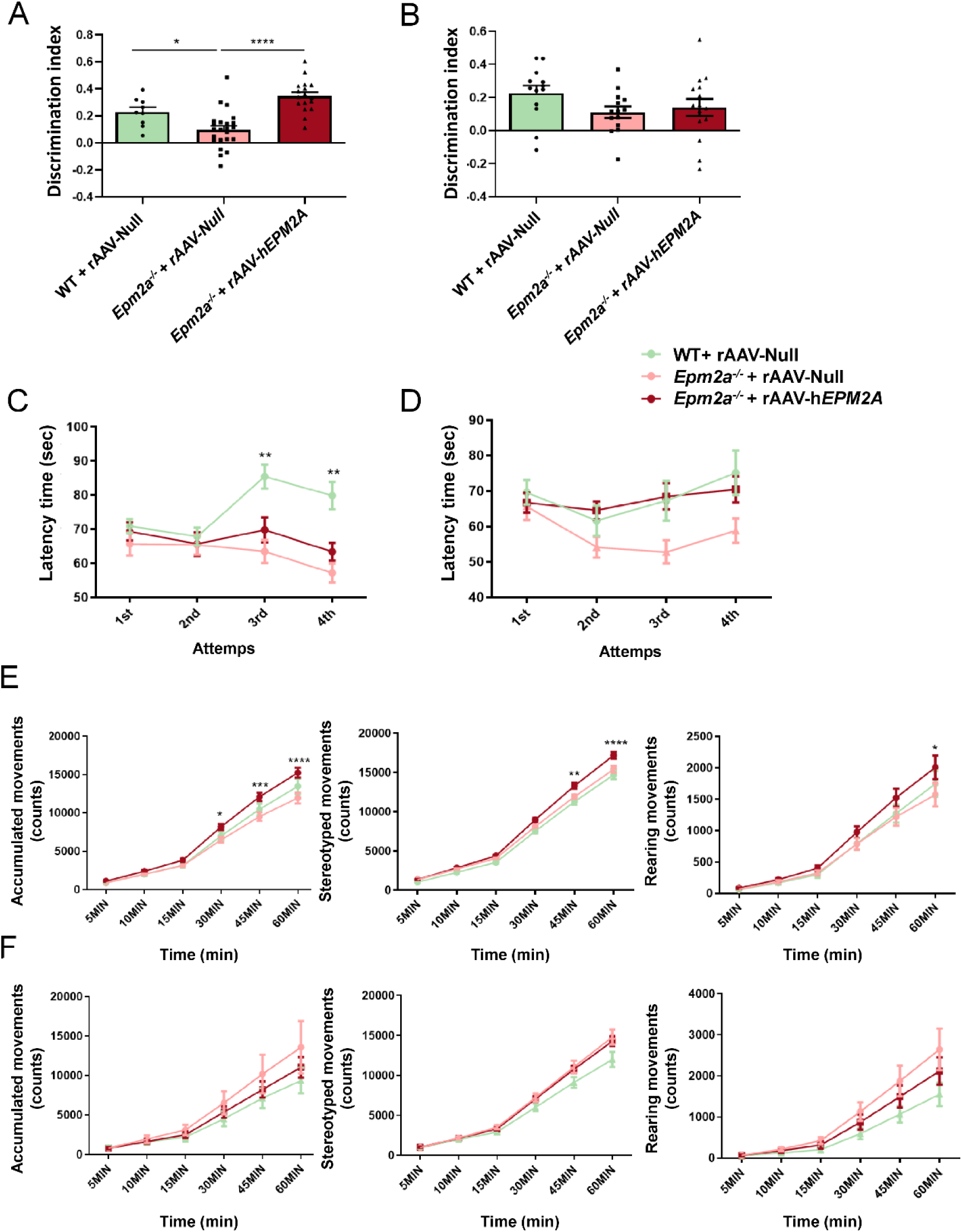
Behavioral studies in *Epm2a^-/-^* mice treated with rAAV-h*EPM2A* or rAAV-Null 3 (A, C, E) and 9 months (B, D, F) after ICV injection. (**A-B**) Memory assessment based on D.I. evaluation in the ORT. (**C-D**) Evaluation of motor coordination based on the latency time to fall from the cylinder of *Epm2a^-/-^* mice. (**E-F**) Analysis of spontaneous accumulated, stereotyped and rearing movements of *Epm2a^-/-^* mice. Data are shown as mean ± SEM. One-way and two-way ANOVA test with Tukey’s multiple comparisons were performed between the experimental groups. * p < 0.05, ** p < 0.01, *** p < 0.001, **** p < 0.0001. n = 15-25 mice per group and experiment

### Treatment with rAAV-h*EPM2A* in *Epm2a^-/-^* mice reduces EEG power, frequency of interictal epileptiform discharges (IEDs), and their heightened sensitivity to PTZ

Video-EEG recordings were performed on 12-month-old mice (9 months after ICV injection) to examine epileptic-like activity. *Epm2a^-/-^* mice treated with rAAV-h*EPM2A* exhibited normalized basal activity rhythms, showing lower beta and gamma wave power spectra compared to *Epm2a^-/-^* mice injected with rAAV-Null, and similar to WT mice **(Fig. 5A)**. Following the administration of a subconvulsive dose of PTZ (30 mg/kg), power spectra decreased in all groups. However, the decrease was more significant in *Epm2a^-/-^* mice injected with rAAV-Null compared to WT or *Epm2a^-/-^* mice treated with rAAV-h*EPM2A*, whose power spectra were similar **(Fig. 5B)**. Additionally, 3- and 9-months post-injection, *Epm2a^-/-^* mice treated with rAAV-h*EPM2A* displayed a reduced incidence of myoclonic jerks in response to IP injection of PTZ (30 mg/kg), compared to *Epm2a^-/-^* mice injected with rAAV-Null **(Fig. 5C-D)**. After IP injection of 50mg/kg PTZ, 12-month-old *Epm2a^-/-^* mice treated with the therapeutic vector exhibited reduced susceptibility to generalized tonic-clonic (GTC) seizures and a lower mortality rate **(Fig. 5E-F).**

**Figure 5.**
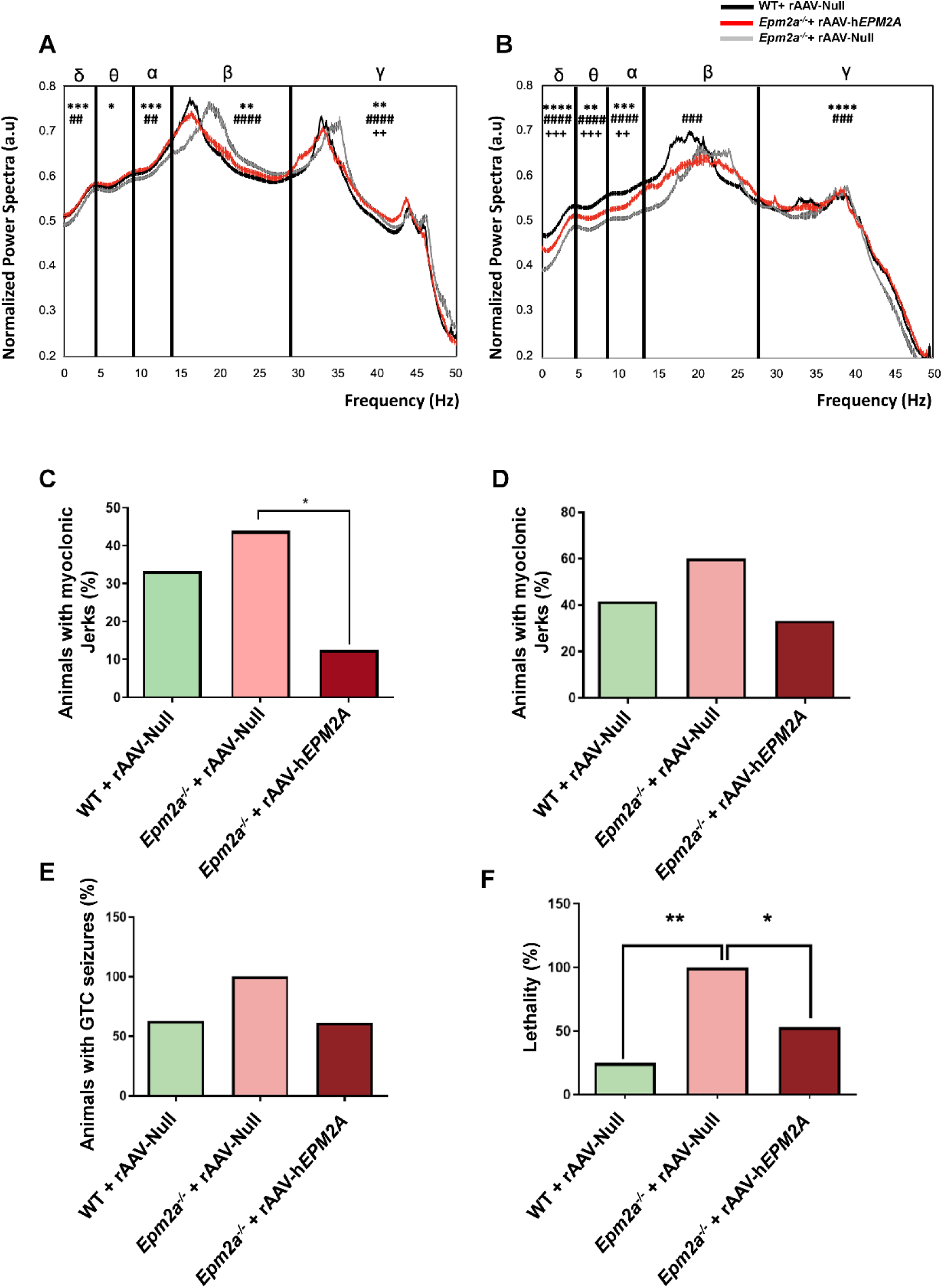
Analysis of EEG and susceptibility to PTZ in WT and *Epm2a^-/-^* mice 3 and 9 months after ICV injection of rAAV-h*EPM2A* or rAAV-Null vectors. (**A**) Representative normalized power spectra obtained from baseline EEG recordings (48 hours). (**B**) Representative normalized power spectra obtained from EEG recordings after IP injection of PTZ at subconvulsive doses (30 mg/kg). Data are shown as mean ± SEM. A one-way ANOVA test with Tukey’s multiple comparisons was performed between the AUC obtained after plotting the EEG powers of all experimental groups. * p < 0.05, ** p < 0.01, *** p < 0.001, **** p < 0.000.1. Symbols indicate: * *Epm2a^-/-^* mice treated with rAAV-h*EPM2A* vs *Epm2a^-/-^* mice injected with rAAV-Null; ^#^ WT injected with rAAV-Null vs *Epm2a^-/-^* mice injected with rAAV-Null; ^+^ WT injected with rAAV-Null vs *Epm2a^-/-^* mice treated with rAAV-h*EPM2A*. n = 2-4 mice per group. (**C-D**) Percentage of animals with myoclonic jerks after IP injection of 30 mg/kg PTZ in WT and *Epm2a^-/-^* mice 3 (**C**) or 9 months (**D**) after ICV injection. (**E**) Percentage of animals with GTC seizures, and (**F**) percentage of lethality after IP injection of 50 mg/kg PTZ in WT and *Epm2a^-/-^* mice 9 months after ICV administration of rAAV-h*EPM2A* or rAAV-Null. Data are shown as percentages. A Fisher’s exact test was performed between the experimental groups. * p < 0.05, ** p < 0.01. n = 15-25 mice per group and experiment

Analysis of spontaneous **(Supplementary Fig. S3A)** and 30 mg/kg PTZ-induced interictal epileptiform discharges (IEDs) **(Supplementary Fig. S3B)** indicated that *Epm2a^−/−^* mice treated with rAAV-h*EPM2A* experienced a reduced number of IEDs than those injected with rAAV-Null, although differences were not statistically significant.

### Expression of human laforin in the brain of *Epm2a^-/-^* mice led to significant changes in critical molecular pathways, as revealed by proteomic and phosphoproteomic analyses of the hippocampus

We performed a proteomic analysis in the hippocampus of 6-month-old *Epm2a^-/-^* mice treated with rAAV-h*EMP2A* compared to *Epm2a^-/-^* and WT mice injected with the rAAV-Null vector. Given the pathophysiological characteristics of Lafora disease, our attention was focused on proteins involved in glycogen metabolism, oxidative stress and regulation of misfolded protein, protein degradation via UPS and neuronal hyperexcitability. Proteins that exhibited a Zq comparison with a p value less than 0.05 (p<0.05) were classified as differentially abundant proteins (DAPs) in each respective comparison.

*Epm2a^-/-^* mice injected with the rAAV-Null vector exhibited increased levels of enzymes involved in glycogen metabolism, including glycogen synthase (GS), glycogenin (GYG), glycogen phosphorylase (PYGB/GP), and glycogen debranching (AGL/GDE), compared to the WT group **(Fig. 6A)**. Expression of human laforin reduced these levels in the hippocampus of *Epm2a^-/-^* treated mice **(Fig. 6A-B).**

**Figure 6.**
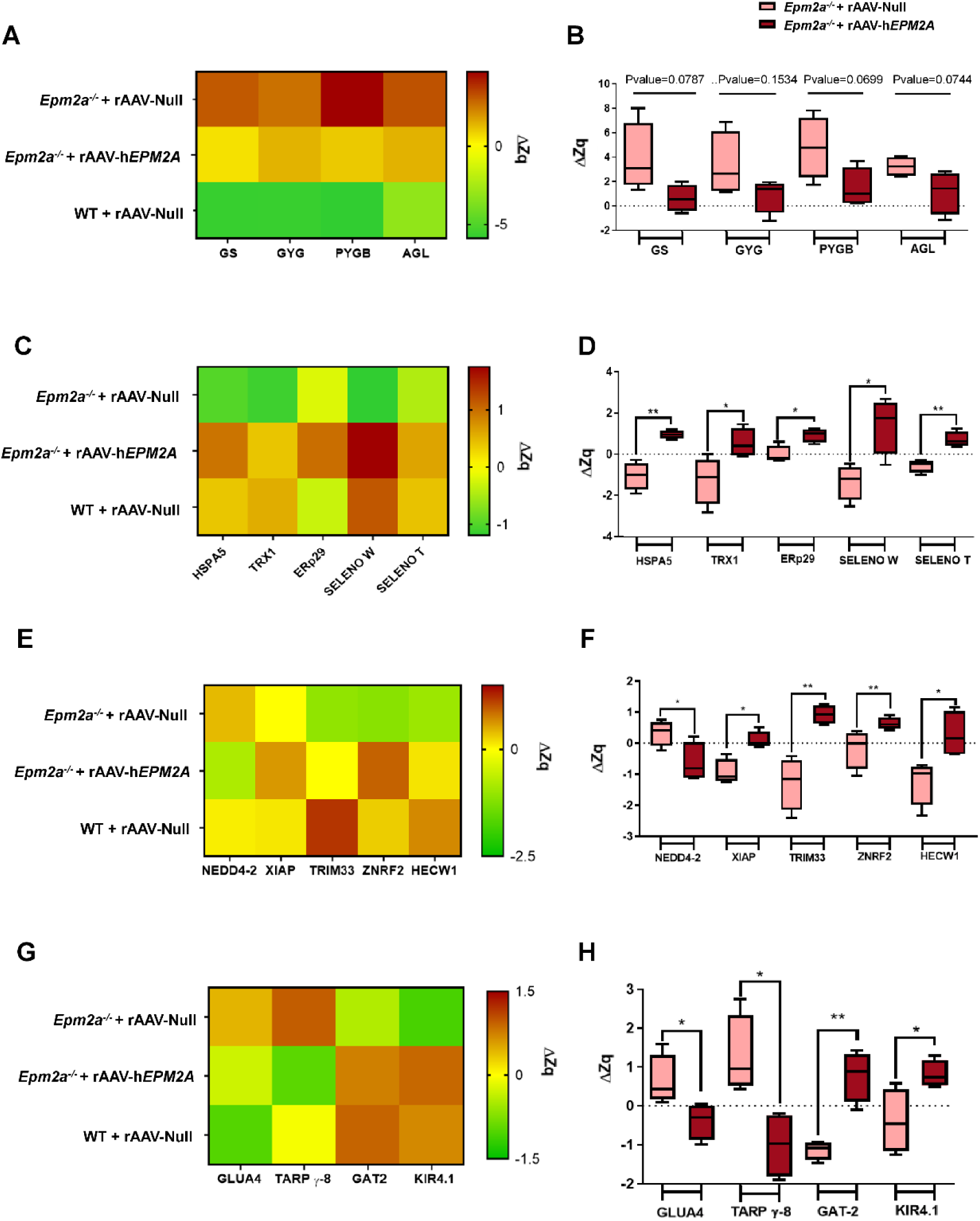
Variations in the abundance of proteins (ΔZq) in the hippocampus of WT and *Epm2a^-/-^* mice 3 months after ICV injection of rAAV-h*EPM2A* or rAAV-Null vectors. (**A-B**) Differences in the abundance of proteins involved in glycogen metabolism. (**C-D**) Variations in the protein abundance related to proteostasis and reactive oxygen species (ROS) regulation. (**E-F**) Differences in the abundance of proteins involved in protein degradation through UPS (**G-H**) Variations in the abundance of proteins associated with neuronal excitability. Data are shown as means of standardized log2 ratio at the protein level (Zq). Whiskers in box plots show the minimum and maximum values. A t-test was carried out between the experimental groups. * p < 0.05, ** p < 0.01. n = 4 mice per group

In *Epm2a^-/-^* mice injected with the rAAV-Null vector, reduced abundance of proteins involved in protein folding and oxidative stress was observed. These proteins included binding immunoglobulin protein chaperone (BiP/HSPA5), thioredoxin (TRX1), endoplasmic reticulum protein 29 (ERp29), selenoprotein W (SELENOW), and selenoprotein T (SELENOT), compared to WT mice **(Fig. 6C)**. Remarkably, the expression of human laforin resulted in increased levels of these proteins **(Fig. 6C-D)**. Next, we explored whether the increased abundance of BiP/HSPA5, a participant in the unfolded protein response (UPR), was a consequence of the abundant release of exogenous laforin, or of a beneficial effect of laforin expression. To address this, we evaluated the interaction between this chaperone and the transmembrane stress sensor proteins inositol-requiring kinase 1 (IRE1) and PKR-related ER kinase (PERK). Coordinated protein response analysis revealed no significant changes in these proteins in 6-month-old *Epm2a^-/-^* mice treated with rAAV-h*EPM2A* compared to *Epm2a^-/-^* mice injected with rAAV-Null vector (**Supplementary Fig. S4A-B)**, suggesting that the increase in proteins related to protein misfolding in the ER is likely a positive response to laforin expression.

*Epm2a^-/-^* mice injected with rAAV-Null also exhibited decreased abundance of several E3 ubiquitin ligases, including tripartite motif-containing 33 (TRIM33), zinc and ring finger 2 (ZNRF2) and HECT, C2 and WW domain containing E3 ubiquitin protein ligase 1 (HECW1), compared to WT mice **(Fig. 6E)**. Treatment with rAAV-h*EPM2A* led to an increased abundance of these proteins in the hippocampus of mice **(Fig. 6E-F)**. Notably, expression of human laforin increased the abundance of X-linked inhibitor of apoptosis (XIAP), an E3 ubiquitin ligase with anti-apoptotic function, and decreased the abundance of NEDD4 like E3 ubiquitin protein ligase (NEDD4.2), whose role in the ubiquitination and endocytosis of GLT-1 in Lafora disease models has been reported **(Fig. 6F).**

In the hippocampus of 6-month-old *Epm2a^-/-^* mice injected with rAAV-Null there was an increased abundance of proteins related to neuronal hyperexcitability **(Fig. 6G)**, including the glutamate ionotropic receptor AMPA type subunit 4 (GLUA4/GRIA4) and calcium voltage-gate channel auxiliary subunit gamma 8 (TARP γ-8). Conversely, there was a reduction in proteins like glial GABA transporter-2 (GAT-2) and potassium inwardly rectifying channel, subfamily J, member 10 (KIR4.1), compared to WT mice **(Fig. 6G)**. Expression of human laforin restored protein levels to those observed in WT mice **(Fig. 6G-H)**.

Finally, we analyzed the levels of peptide phosphorylation in the hippocampus of 6-month-old *Epm2a^-/-^* mice treated with rAAV-h*EMP2A* compared to *Epm2a^-/-^* and WT mice injected with the rAAV-Null vector. The abundance of phosphorylated/dephosphorylated peptides was assessed using the standardized log2 ratio, Zp, for several proteins relevant to multiple molecular pathways in Lafora disease **(Table 1)**. These included enzymes related to glycogen metabolism, such as GS, 1,4-alpha-glucan branching enzyme (GBE1) and PYGB/GP, many kinases and phosphatases, such as RAC-beta serine/threonine-protein kinase (AKT2), AKT3, various PKC isoforms, phosphatidylinositol 3,4,5-trisphosphate 3-phosphatase and dual-specificity protein phosphatase PTEN, kinase suppressor of RAS 1 (KSR1), and mitogen-activated protein kinase 7 (MAPK7). Differential phosphorylation was also observed in neurotransmitter receptors and channels, such as ionotropic glutamate receptor, NMDA2A, glutamate receptor (GRIA1), isoform 4 of glutamate receptor 2 (GRIA2), potassium voltage-gated channel subfamily KQT member 2 (KCNQ2), KCNQ3, and disc larg homolog 1 or synapse-associated protein 97 (DLG1/SAP97) **(Table 1)**, all of which may be involved in the molecular mechanisms underlying Lafora disease.

**Table 1.**
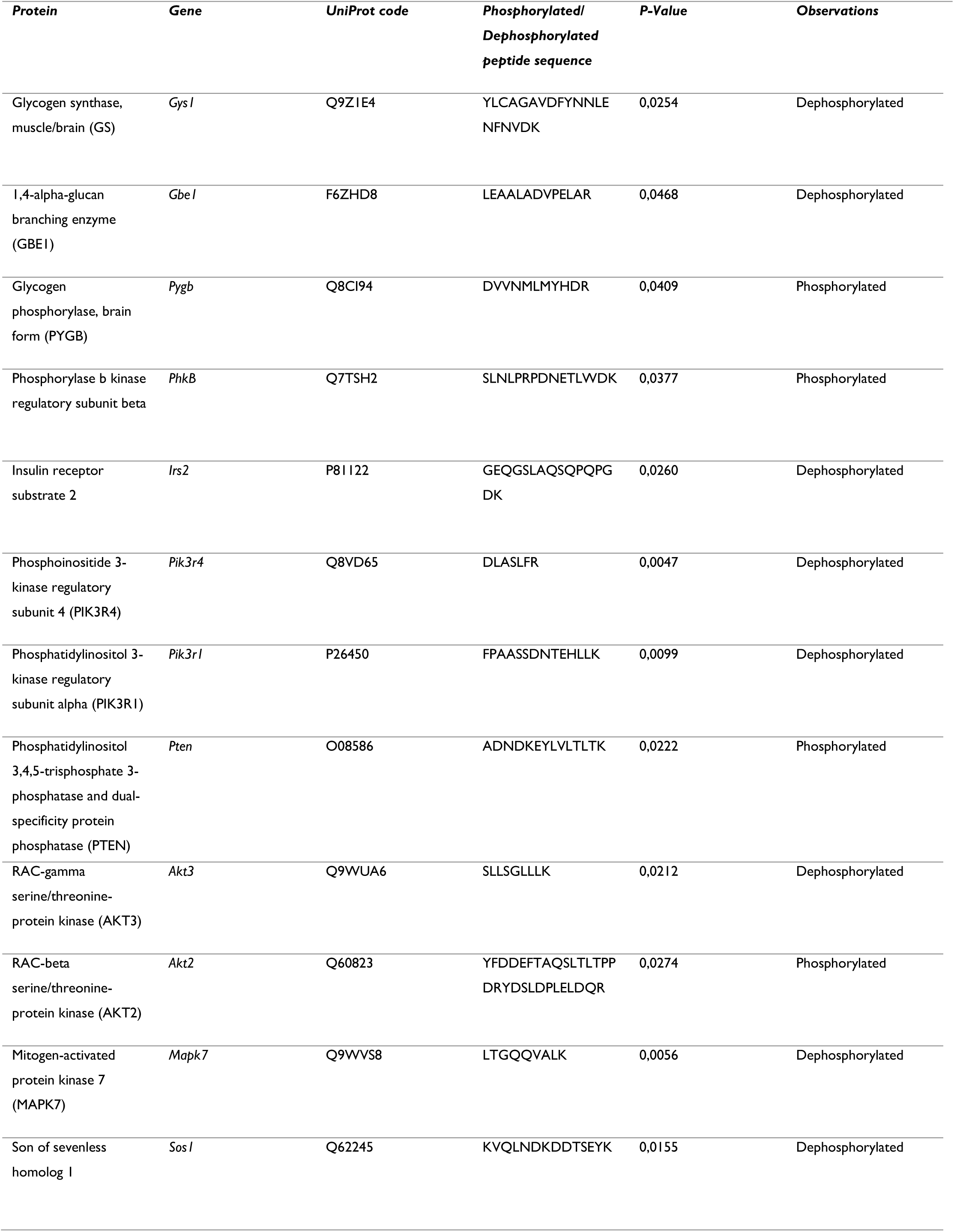
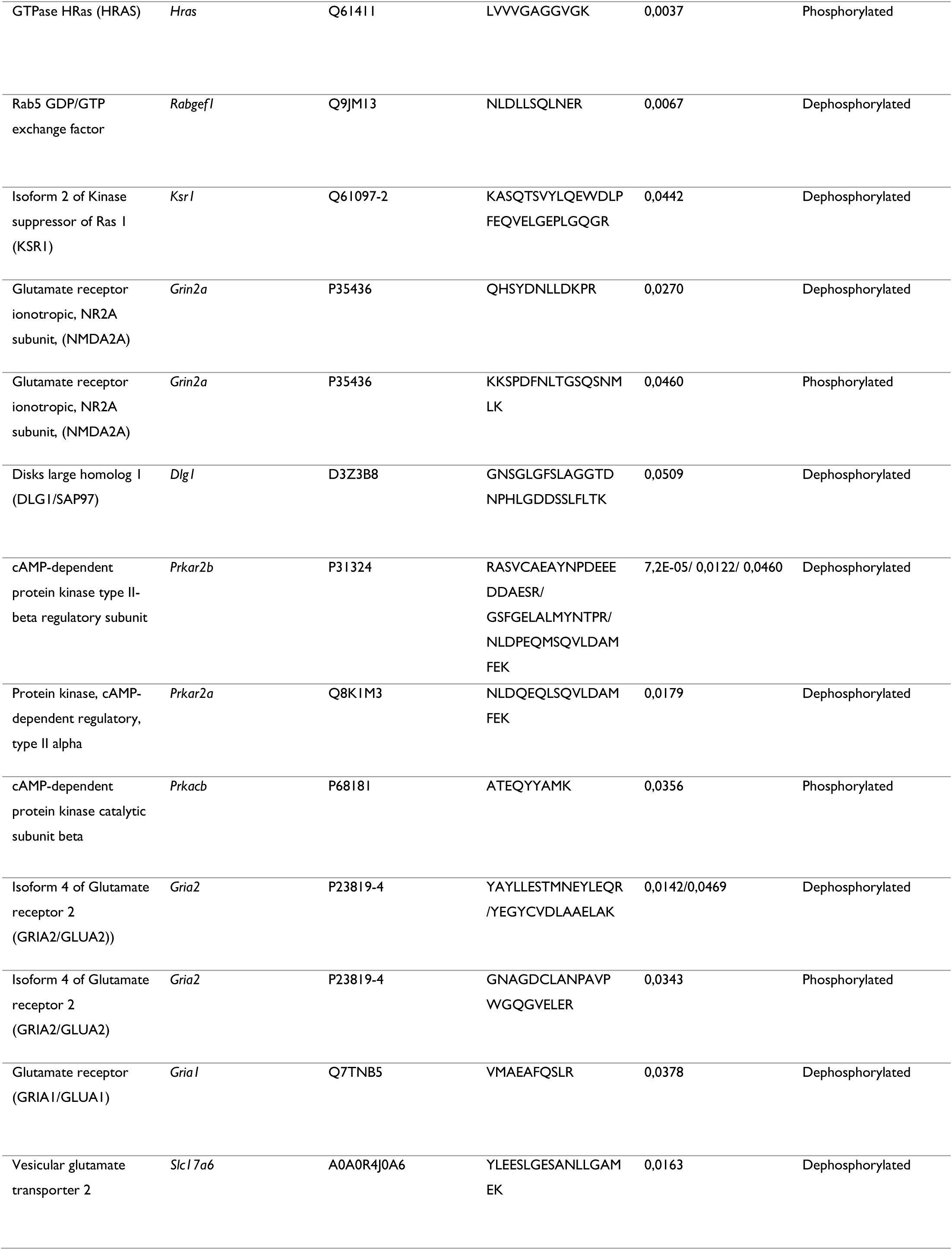
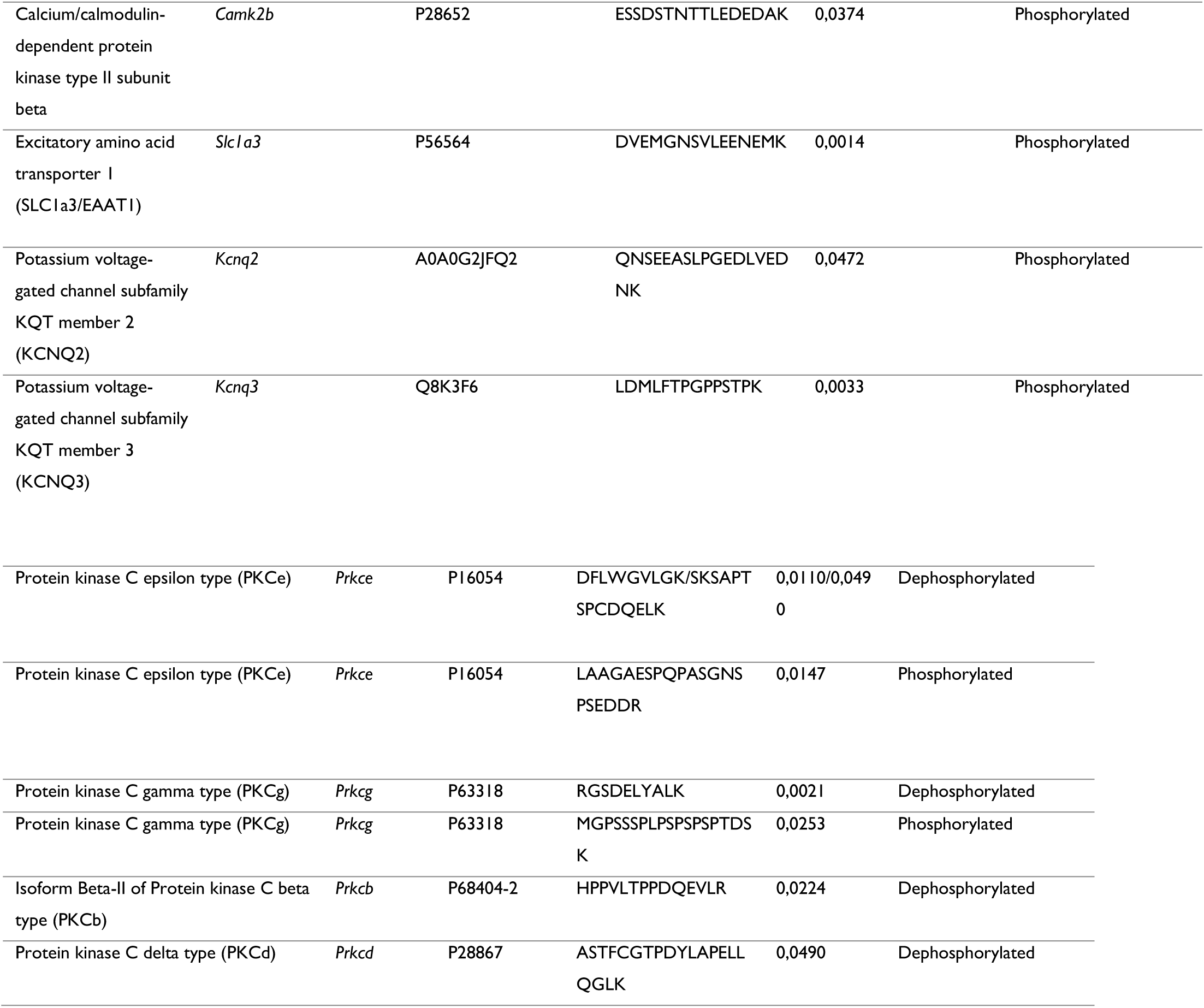
Phosphorylation/dephosphorylation profile of peptides related to Lafora disease in *Epm2a^-/-^* mice treated with rAAV-h*EPM2A*compared to *Epm2a^-/-^* mice injected with rAAV-Null. *P-*value indicates the significant differences between the means of Zp (ΔZp)

## Discussion

This study demonstrates the significant therapeutic impact of gene replacement in a mouse model of Lafora disease. This is the first-ever example of a successful gene therapy with enduring benefits in a preclinical mouse model of Lafora disease. We developed a gene replacement therapy with a rAAV2/9 vector to restore the expression of the *EPM2A* gene and recover the function of laforin in the *Epm2a^-/-^* mouse model of Lafora disease. Evaluation of the effects of this form of therapy involved testing memory abilities, motor coordination, spontaneous locomotor activity, epileptic activity and neuronal hyperexcitability, along with neuropathological and molecular changes. Treatment with the rAAV-h*EPM2A* vector improved many neurological outcomes 3 and 9 months after a single ICV injection. Proteomic and phosphoproteomic analysis revealed changes in important molecular signaling pathways associated with the expression of human laforin.

Disruption of either laforin or malin results in the formation of aggregates of polyglucosans^14–17^. In our study, we observed a significant reduction in LB formation in *Epm2a^-/-^*mice treated with rAAV-h*EPM2A*, as compared to mice injected with rAAV-Null. The laforin-malin complex controls glycogen synthesis by causing the proteasome-dependent degradation of GS, GDE, and PTG^11,12^. Proteomic analysis revealed that untreated *Epm2a^-/-^* mice exhibited an abnormal abundance of these enzymes involved in glycogen metabolism in the hippocampus compared to WT mice. Gene replacement therapy reversed this enzymatic imbalance, restoring enzyme levels to those observed in WT mice, thus correcting abnormal glycogen accumulation and LB formation.

Alteration in protein homeostasis, ROS regulation, glutamate homeostasis, or UPS are the main pathophysiological conditions in most neurodegenerative disorders, including Lafora disease. The laforin-malin complex contributes to mitigating the toxicity caused by misfolded proteins^13^ via the UPS. Moreover, LBs sequester a variety of proteins involved in proteostasis and autophagy^22,48,49^, highlighting the significance of protein homeostasis and deregulation of ROS in the pathophysiology of the disease^18,19^. These alterations contribute to a significant reactive astrogliosis and neuroinflammatory responses in the brain. Notably, gene replacement therapy reduced neuroinflammation in *Epm2a^-/-^* mice. Chaperones^25,50^, antioxidants^25,27^, and autophagy stimulators^24,25,51^ are examples of compounds that have shown promise in lowering or rescuing neuroinflammation in Lafora disease mouse models. In this regard, our proteomic analysis revealed a substantial increase in proteins associated with protein misfolding, UPS and the regulation of ROS in *Epm2a^-/-^* treated mice. Thus, abundance of TRX1, a proteasome related protein that is decreased in fibroblasts from Lafora disease patients^52^, and BiP chaperone^19^ are increased in *Epm2a^-/-^* mice treated with rAAV-h*EPM2A*. The ERp29 chaperone and the antioxidant selenoproteins T and W are also more abundant in *Epm2a^-/-^* treated mice. Furthermore, rAAV-h*EPM2A* increased the abundance of several E3 ubiquitin ligases. However, NEDD4-2 ubiquitin ligase, which has been previously reported to induce the ubiquitination and endocytosis of GLT-1 in Lafora disease models^53^, exhibits lower abundance. The reduction in the abundance of this protein produces decreased ubiquitination of GLT-1 increasing the levels of the glutamate transporter at the plasma membrane and the glutamate uptake capacity^53,54^. These data strongly suggest that the expression of laforin in treated mice improves neurological functions by reducing misfolded proteins, stimulating UPS and glutamate reuptake and alleviating alterations in ROS, therefore decreasing neurological alterations as memory abilities and motor spontaneous activity and coordination.

Mice treated with rAAV-h*EPM2A* showed fewer myoclonic jerks, GTC seizures, and a lower mortality rate after IP administration of both subconvulsive and convulsive PTZ doses. Furthermore, video-EEG analysis revealed that *Epm2a^-/-^* treated mice exhibited a restoration of normal power spectra, particularly in the gamma range, that is known to be associated with epileptic activity ^55–57^. *Epm2a^-/-^* mice injected with rAAV-h*EPM2A* displayed less decrease in EEG power after PTZ administration across high-frequency waves, indicating decreased PTZ susceptibility^58^. In addition to these findings, other changes consistent with EEG and PTZ sensitivity alterations were noted in proteomic and phophoproteomic studies. A higher abundance of the GABA transporter GAT-2 was observed in the hippocampus of mice treated with the therapeutic vector. Reduced GAT-2 levels have been associated with temporal lobe epilepsy ^59^. This GABA transporter may contribute to reducing neuronal hyperexcitability in treated mice through glutamate-induced GABA release ^60–62^. Phosphorylation of NMDA modulatory subunits (NR1, NR2A and NR2B), and AMPA receptors regulates neuronal excitability, among other neuronal pathways. In the hippocampus of mice treated with rAAV-h*EPM2A*, we observed significant dephosphorylation of NR2A and AMPAR subunits GLUA1 and GLUA2, along with a notable decrease in GLUA4 abundance. These findings suggest that gene therapy with rAAV-h*EPM2A* may decrease neuronal hyperexcitability by reducing glutamatergic transmission.

In addition, we observed a reduction in neuronal hyperexcitability resulting from molecular pathways beyond the glutamatergic and GABAergic systems. Potassium channels are crucial for maintaining the balance of brain excitability. Thus, mice treated with rAAV-h*EPM2A* exhibited restoration of normal KIR4.1 channel levels. Additionally, mutations in KCNQ2 and KCNQ3 are linked to Benign Familial Neonatal Convulsions (BFNC), and phosphorylation of these channels by cAMP-dependent PKA stimulates their activity, reducing epileptic activity ^63,64^. Phosphorylation patterns suggest increased PKA activity in mice treated with therapeutic vectors, with higher phosphorylation of KCNQ2 and KCNQ3 in the hippocampus. These findings may collectively explain the decrease in epileptic activity in *Epm2a^-/-^* mice treated with rAAV-h*EPM2A*.

In conclusion, our study demonstrated that gene replacement therapy with the human *EPM2A* gene resulted in significant neurological improvements in a mouse model of Lafora disease. Treatment at an early symptomatic stage markedly reduced neuroinflammation and LB formation, delayed memory and motor alterations, improved motor coordination, and reduced epileptic activity. Furthermore, proteomic and phosphoproteomic analyses shed light on various mechanisms through which laforin may be inducing these improvements, indicating potential new targets in Lafora disease. Our results with gene replacement therapy open a new avenue for treating this devastating disease.

## Acknowledgments

We thank Miguel Chillón Rodríguez (Viral Vector Production Unit or UPV-UAB-VHIR) and Juan Antonio López del Olmo (CNIC Proteomics Unit) for their technical advice. We also thank the Animal Facility of Instituto de Investigación Sanitaria-Fundación Jiménez Díaz, for their technical assistance.

