## Supplementary Material for "Gene replacement therapy for Lafora disease in the *Epm2a^-/-^* mouse model"

### Supplementary Material and Methods

#### Immunofluorescence, immunohistochemistry and PAS-diastase staining

Mice were anesthetized, transcardially perfused with 4% phosphate-buffered paraformaldehyde, and their brains were removed, dehydrated, and paraffin-embedded. Blocks were then sectioned into consecutive 5-µm-thick sections. For PAS-diastase (PAS-D) staining, sections were rehydrated using decreasing graded alcohols, treated with porcine pancreas α-amylase (5 mg/ml in dH2O) (Merck, Darmstadt, Germany) and processed with the PAS Kit (Merck, Darmstadt, Germany). Following this, they were counterstained with Gill No. 3 hematoxylin (Merck, Darmstadt, Germany). For immunohistochemistry (IHC) and immunofluorescence (IF-P), rehydrated sections underwent boiling in 0.1 M sodium citrate buffer, pH 6.0. Samples were then incubated in blocking buffer (1% bovine serum albumin, 5% fetal bovine serum, 2% Triton X-100, diluted in PBS) and then with primary antibodies diluted in blocking buffer. For IHC, the primary antibodies used were green fluorescent protein (GFP) (1:100 dilution; Abcam, Cambridge, UK; Cat. #ab183734), neuronal nuclei (NeuN) (1:100 dilution; Millipore, Temecula, CA, USA; Cat. # MAB377) and laforin (3µg/mL; Lifespan Biosciences, Washington, US, Cat. #LS B6474). Sections were stained with the Vectastain ABC kit (Vector Laboratories, Burlingame, CA, USA). Immunoreactivity was revealed using diaminobenzidine (Dako Cytomation, CA, USA) and H2O2 and counterstained with Carazzi hematoxylin (Panreac Quimica, Barcelona, Spain). For IF-P, the primary antibody was glial fibrillary acidic protein (GFAP) (1:1000 dilution; Millipore, Temecula, CA, USA; Cat. #MAB360). Secondary antibodies were conjugated to Alexa Fluor 594 (donkey anti-mouse, 1:400 dilution; Abcam, Cambridge, UK, Cat. #ab150108). Samples from 4-6 mice per group were used, and two consecutive sections per animal were stained and analyzed. Images from the same area of the CA1 region of each hippocampus were acquired using a Leica DMLB 2 microscope (Leica, Wetzlar, Germany) connected to a Leica DFC320 FireWire digital microscope camera (Leica, Wetzlar, Germany) for IHC processed sections, and with a Zeiss Axioscope 5 (Zeiss, Jena, Germany) connected to an Axiocam 208 color camera (Zeiss, Jena, Germany) for sections processed via IF-P. Subsequently, LBs and NeuN- or GFAP- positive cells were quantified by two researchers using ImageJ software (NIH, Bethesda, MD, USA). Reported values represents the average of these quantifications.

#### Spontaneous locomotor activity

Spontaneous movements were monitored with a computerized actimeter (Harvard Apparatus, Holliston, MA, USA) that recorded the number of times each mouse crosses the open field through infrared light beam breaks. The SEDACOM 1.4 software (Harvard Apparatus, Holliston, MA, USA) was utilized to analyze the spontaneous, rearing, and stereotyped movements at 5, 10, 15, 30, 45, and 60-minute intervals.

#### Motor coordination

Motor coordination and balance were assessed using the rotarod test (Harvard Apparatus, Holliston, MA, USA). The mice underwent two days of training. On the first day, mice were placed on the rotarod at a constant speed of 4 rpm for 60 seconds. On the second day 2, they were trained with speed increasing from 4 to 8 rpm. In subsequent test sessions over two days, only mice able to stay on the rod for 60 seconds were included to minimize learning related variations. The latency time to fall from the cylinder was recorded during two sessions each day, with speed increasing from 4 to 40 rpm, and a maximum time limit of 5 min.

#### Object recognition task

The object recognition task (ORT) was used to assess episodic memory retention. Mice were individually familiarized with a dark box in an open field for 10 min. Two hours later, two identical objects (A and B) were placed in the center of the box. Each mouse had 10 min to freely explore the objects with exploration times (tA and tB) recorded. After two hours, a new object C replaced object B, and the exploration times (tA and tC) were measured. Virtual timers generated by the XNote Stopwatch software were used to measure explorations times when mice examined objects from 2 cm or less. A discrimination index (D.I.) was calculated using the following equation: D.I. = (tC-tA) / (tC + tA).

#### Sensitivity to Pentylenetetrazole (PTZ)

To analyze neuronal hyperexcitability, PTZ was administered via intraperitoneal injection, using two doses: 30 mg/kg (subconvulsive dose, rarely causing generalized tonic-clonic seizures in WT animals) and 50 mg/kg (convulsive dose). The subconvulsive dose was used to determine the percentage of mice displaying myoclonic jerks. The convulsive dose was administered to evaluate the percentage of animals experiencing GTC seizures, their duration, the time to the first myoclonic or GTC seizure, and the lethality. Each animal was observed for 45 minutes by two researchers.

#### Video-EEG analysis

A plastic pedestal (Plastics1, Virginia, USA) with trimmed electrodes was surgically implanted and secured with acrylic resin onto the skull. Post-surgical pain was managed with meloxicam (5 mg/kg) (Boehringer Ingelheim, GA, USA). Animals were allowed one week for recovery before testing. Video-EEG recordings were obtained using a wireless transmitter (Epoch, CA, USA) attached to the pedestal, and the data were digitally recorded on a computer under free-motion conditions. Mice were observed in their home-cages under basal conditions and, after 48 hours, recorded for 30 minutes following PTZ injections. The sampling rate was 250 Hz and a 50 Hz notch filter was applied. Mouse behavior was captured using digital video cameras. EEG data were analyzed automatically and manually using the Acknowledge® 5.0 software (Epoch, CA, USA) excluding periods with signal loss or artifacts. After applying the Comb Band Stop filter and a Blackman window, power spectra were calculated using a spectral estimator based on autoregressive processes ensuring normalized amplitudes for consistent peak-to-peak ranges. An automated seizure analysis was also conducted, closely monitoring and thoroughly analyzing both seizures and IEDs (interictal epileptiform discharges).

### Supplementary figures

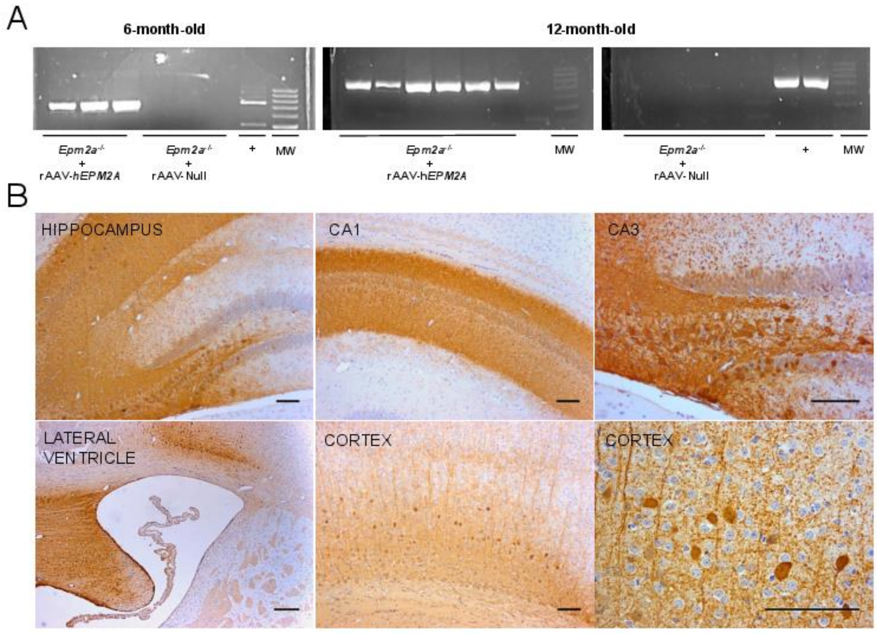


**Supplementary Figure 1. Transduction of the rAAV-*GFP* and rAAV-h*EPM2A* vector into CNS cells.** (A) RT-PCR shows efficient transcription of laforin mRNA in brain of mice treated with the vector. (B) IHC with an anti-GFP antibody revealing the ability of the vector to transduce CNS cells across the hippocampus, cortex, and periventricular areas. n= 4-6 mice per group and experiment. Scale bar =100µm

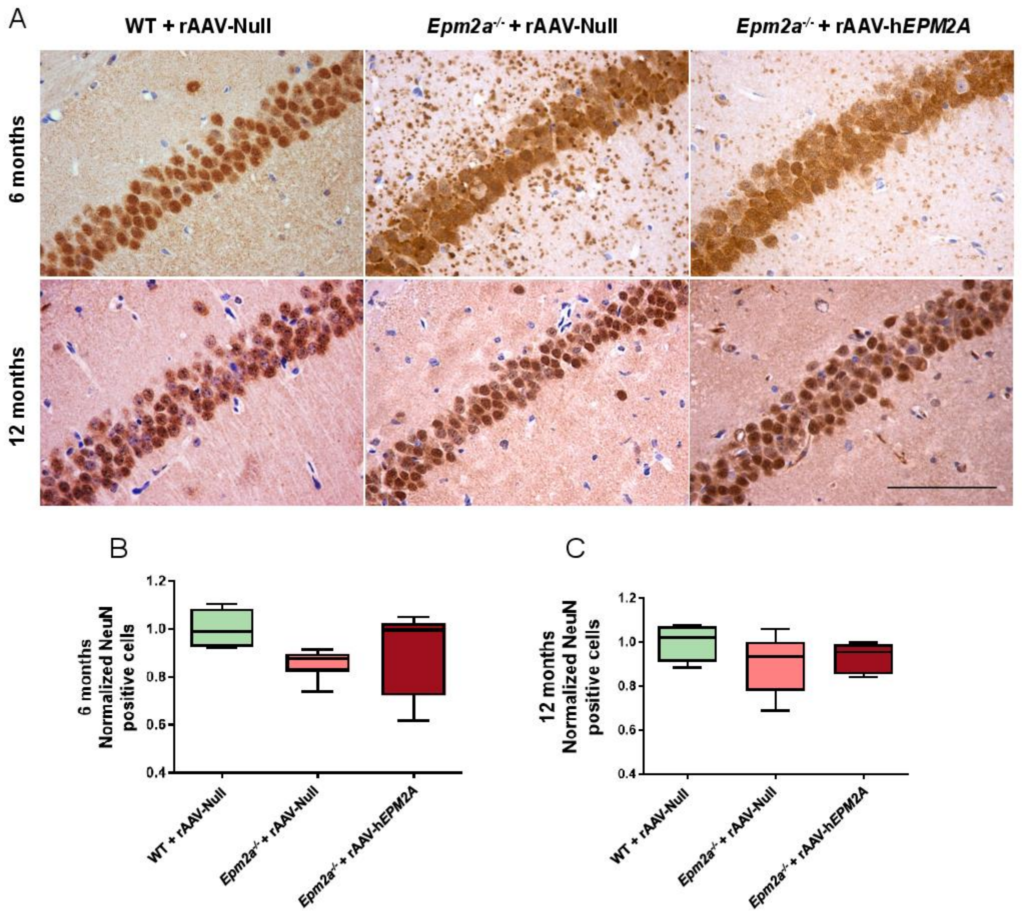


**Supplementary Figure S2. Effects of rAAV-h*EPM2A* treatment on neurodegeneration in the CA1 region of the hippocampus of *Epm2a*$^{-/-}$ mice using a NeuN antibody**. (A) Images of NeuN immunostaining in the hippocampal CA1 region at 6 and 12 months, 3 and 9 months after rAAV-h*EPM2A* injection. Quantification of NeuN-positive cells 3 (B) and 9 months (C) after treatment. A non-parametric Kruskal-Wallis test was performed followed by Dunn´s multiple comparisons. Results are expressed as median of independent samples. Whiskers in box plots show the minimum and maximum values. No significant differences were found between groups. n= 4-6 mice per group and experiment. Scale bar =100 µm

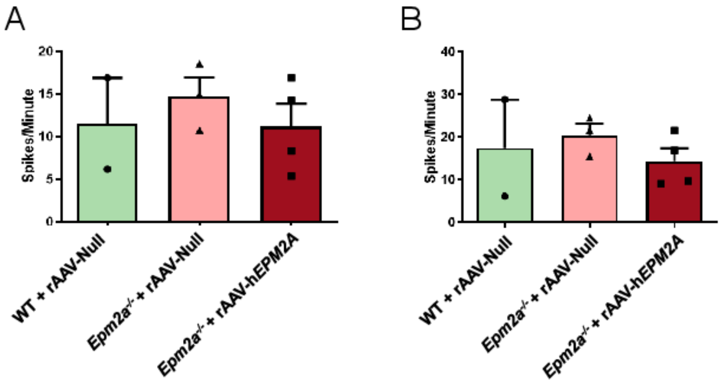


**Supplementary Figure S3. Analysis of spontaneous and PTZ-induced interictal epileptiform discharges (IEDs)**. A tendency to present more IEDs was observed in *Epm2a*$^{-/-}$ mice injected with rAAV-Null compared to *Epm2a*$^{-/-}$ mice treated with rAAV-h*EPM2A* or to WT in basal (A) or PTZ-induced (B) condition 9 months after ICV injection. One-way ANOVA followed by Tukey's multiple comparisons test was performed between the experimental groups. No statistical differences were found. n= 2-4 mice per group and experiment

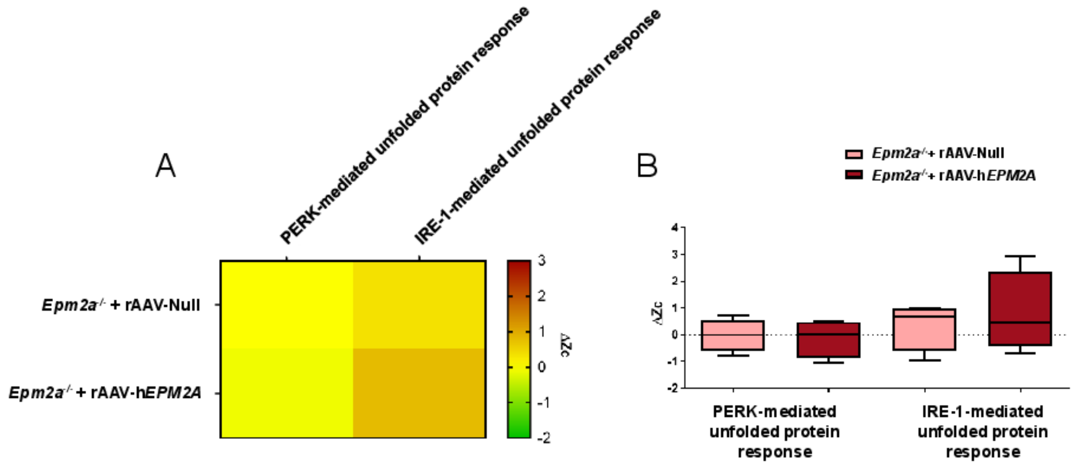


**Supplementary Figure S4. Coordinated protein response analysis to assess the potential interaction between the chaperone BiP/Hspa5 and IRE1 or PERK**. (A) Representative heatmap showing no significant changes in these pathways in 6-month-old *Epm2a*$^{-/-}$ mice treated with rAAV- h*EPM2A* compared to those injected with rAAV-Null vector. (B) Results expressed as comparative means of standardized log2 ratio of categories levels (Zc). Whiskers in box plots show the minimum and maximum values. n = 4 mice per group
